# A bio-orthogonal and covalent 5 kDa small protein tag

**DOI:** 10.64898/2026.04.10.717487

**Authors:** Ulrich Pabst, Jana Rossius, Christina Holmboe Olesen, Melissa Birol, Johannes Broichhagen

## Abstract

Precise and minimally perturbative protein labelling remains a key challenge for studying biomolecular function in living systems. Here, a minimal way to specifically label proteins based on SNAP-tag and intein-mediated protein splicing reaction is introduced. Termed CLUSTER (for <u>C</u>hemical <u>L</u>abel-<u>U</u>nfold-<u>S</u>plice <u>T</u>echnology <u>E</u>nables <u>R</u>ecombination), this chimeric platform supports efficient labelling across diverse targets in living cells by retaining a fluorescent, 5 kDa sized peptide on a fusion protein of interest after splicing. A bacterial screening workflow was developed to optimize the reaction efficiency and construct design. Quantitative characterization using fluorescence polarization provides mechanistic insight into labelling efficiency and dynamics, while molecular dynamics simulations elucidate its stability, grasping the intricate nature of protein behaviour upon covalent labelling. This bio-orthogonal labelling technology allows for a versatile and minimally invasive approach for protein labelling, providing a powerful tool to probe protein behavior in native cellular systems.

## INTRODUCTION

Fluorescence imaging of proteins of interests (POIs) requires specific labeling with chromophores, often achieved via immunohistochemistry or as fusion proteins.^1–3^ However, this adds considerable size to the POI, with the disadvantage that it may perturb the proteins native function and behaviour. Antibodies comprise large molecular entities (150 kDa, **Fig. 1A**), substantially exceeding the size of, for instance, their epitopes (HA, 3 kDa), as well as many, transmembrane receptors like GLP1R (55 kDa) and its endogenous ligand GLP1 (4 kDa). Alternatively, genetic fusion to fluorescent proteins (FPs, ∼27 kDa) are widely used. However, FPs are inferior in their photophysical properties to organic fluorophores that display augmented brightness and photostability, and which can be specifically and rapidly reacted with self-labeling protein tags (SLPs),^4^ like the HaloTag Protein (HTP, 33 kDa) and SNAP-tag (20 kDa). Still comparably large in size, folded tags, however, may disrupt conformations, functions and dynamics of POIs.^5,6^ This has been addressed by for instance split enzyme complementation, which has led to improved knock-in strategies^7^ and building protein based sensors^8^. Among the pi-clamp system (0.5 kDa) that uses sticky fluorinated substrates^9^ that may not be ideal for live cell applications, small covalent tagging systems remain scarce.

**Figure 1.**
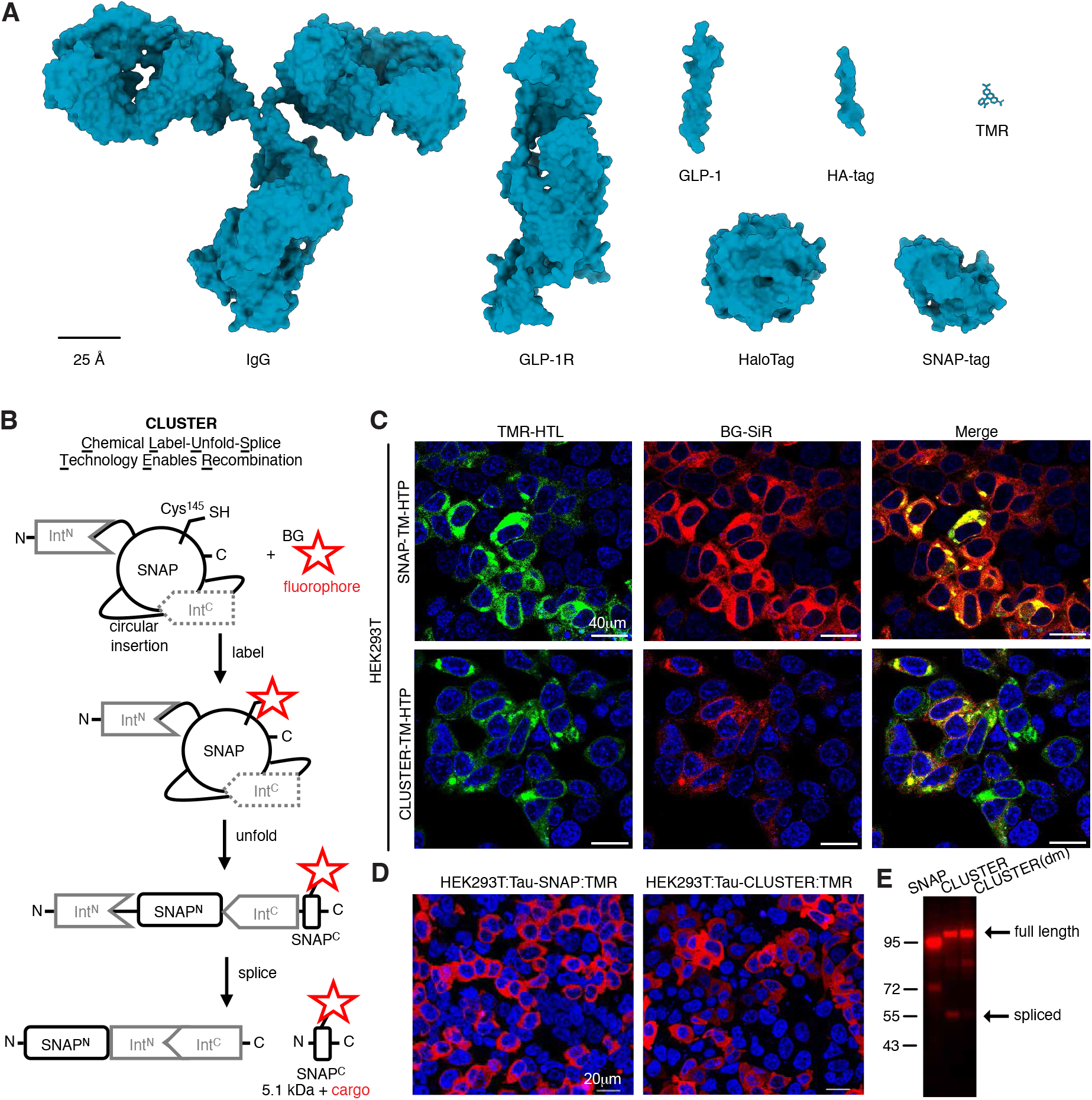
Protein sizes, CLUSTER design and expression. **A)** Size comparison of various relevant molecular structures, including immunoglobulin, GLP-1 and GLP-1R, various tags (HaloTag, SNAP-tag, HA-tag), and tetramethylrhodamine (TMR). **B)** SNAP-tag is endowed *N*-terminally with gp41-1^N^ and inserted with gp41-1^C^ between position 132-133, upstream of the reactive residue Cys145. Upon SNAP-tag reaction with its BG substrate, the construct becomes less stable and the inteins assemble for splicing, transferring the labelled *C*-terminal SNAP residue onto a *N*-terminal protein of interest. **C)** Live HEK293T cells transiently transfected with SNAP–TM–HTP or CLUSTER–TM–HTP, stained with TMR-d12-HTL and BG-SiR-d12. **D)** Live HEK293T cells transiently transfected with Tau–SNAP or Tau– CLUSTER, stained with TMR-d12-HTL and BG-SiR-d12. **E)** SDS-PAGE of protein extract from (D), including a dead mutant (dm), showing only spliced and fluorescently labelled product for Tau-CLUSTER.

### RESULTS

Herein, we describe a conceptually new protein-labelling strategy that 1) minimizes tag-size, is 2) controllable in space and time, is 3) bio-orthogonal, and as such 4) applicable in live cells and 5) uses synthetic, bright fluorophores. The approach utilizes the SNAP-tag, which reacts with benzylguanine (BG) substrates, forming a covalent bond on Cys145.^10,11^ We inserted the *C*-terminal domain of a *trans*-splicing split intein (gp-41-1^C^)^12^ between positions 132/133, just upstream of the reactive cysteine residue (**Fig. 1B**). Upon SNAP-labelling, gp-41-1^C^ will become accessible to its natural interaction partner, gp-41-1^N^, that resides on the *N*-terminus of SNAP. When the stoichiometric split parts meet, they splice the fluorophore containing peptide (47 amino acids, i.e., 5.1 kDa, excluding cargo) onto the POI.

This engineered system is termed <u>CLUSTER</u>, standing for <u>C</u>hemical <u>L</u>abel-<u>U</u>nfold-<u>S</u>plice <u>T</u>echnology <u>E</u>nables <u>R</u>ecombination. We subcloned the first design (CLUSTER_238_) replacing the SNAP-tag in our reported SNAP-TM-HTP plasmid^13^ (TM = transmembrane domain). We first transfected HEK293T cells with the parent SNAP-TM-HTP and observed clean HTP and SNAP labelling using bright TMR-d12-HTL (HTL = HaloTag Ligand) and BG-SiR-d12 (ref^14^) (BG = *O*^6^-benyzlgunaine, i.e. SNAP-tag substrate), respectively (**Fig 1C**). Substitution with CLUSTER_238_–TM–HTP in a similar experiment, yielded detectable dual TMR and SiR signals, albeit at reduced intensity. To assess performance of CLUSTER on intrinsically disordered proteins, we next examined the Tau protein as a benchmark, a microtubule-associated protein implicated in Alzheimer’s Disease and tauopathies^15^, which shows complex behaviour such as phase separation^16^. We transfected HEK293T cells with Tau-SNAP or Tau-CLUSTER_238_ and labelled both samples successfully with BG-TMR-d12 for subsequent confocal microscopy (**Fig 1D**). While fluorescence signals were recorded for both instances, we harvested the cells and loaded them on SDS-PAGE, for which a spliced product was detectable exclusively for Tau-CLUSTER_238_, with an intein-dead mutant (DM) serving as control (**Fig. 1E**, and see SI). This demonstrates that 1) CLUSTER can be be expressed in mammalian system and 2) being labelled with SNAP-tag substrates, both 3) intra- and extracellularly, and 4) is amenable to intein-driven splicing after it has reacted with a substrate.

We next created a construct (CLUSTER_238_–HTP) (**Fig. 2A**) as a model system for screening in *E. coli*. BG-SiR-d12 and TMR-d12-HTL were used for labelling SNAP and HTP, respectively, the latter for expression control and normalization before cells were lysed and subjected to SDS-PAGE. We successfully observed tagged proteins under a fluorescent scanner, both spliced (lower band) and non-spliced (upper band) (**Fig. 2B**). Encouraged by this, we performed a glycine linker deletion screen (i.e., G95, G96, G232, G270 and combinations thereof) that reside as linkers for gp-41-1^N^ (G95/96) and flank the inserted gp-41-1^C^ (G232/270) to find mutant CLUSTER_277_ working binary, i.e. where SiR-signals are only detectable at the lower molecular weight band (**Fig. 2B**). This indicates that labelling occurs on the full-length protein before splicing. To gain a deeper insight into the mechanism, we performed kinetic measurements by fluorescence polarization on recombinantly expressed SNAP, CLUSTER_277_–HTP and SNAP^N^–gp-41-1^C^–SNAP^C^ (i.e. CLUSTER_340_) ± gp41-1^N^–HTP and when exposed to BG-TMR-d12. While SNAP showed kinetics towards full labelling within minutes (*k*_obs_ = 22.2 mHz) (**Fig. 2C**), CLUSTER_277_–HTP gave a rise in polarization (*k*_obs_ = 0.932 mHz), after which a drop occurred, displaying complex parallel reactions of labelling and splicing (**Fig. 2D**). To gain a better understanding of this complexity, we tested CLUSTER_340_ that showed a steady increase in polarization due to covalent reaction, however, with slower kinetics (*k*_obs_ = 0.721 mHz), reaching a plateau after approximately 2 hours (**Fig. 2E**). As the signal remained stable, we added gp41-1^N^–HTP, which lead to a decrease in polarization indicating 1) successful splicing and, similarly important, 2) dropping polarization values to approximately 55 mP. The drop in polarization can be attributed to the fluorophore’s more disordered environment with several more degrees of freedom compared to when bound to folded structures. Importantly, BG-TMR-d12 was unable to react with only the *C*-terminal SNAP^C^, as demonstrated by orthogonal control reactions of SNAP^C^ (synthesized by SPPS). This was achieved with maleimide-linked TMR-d12 (for cysteine reaction) and BG-TMR-d12 (that does not react) and subsequent fluorescence polarization readouts, where the strongly significant observed signal difference to the more reactive maleimide species indicated no significant labelling reactivity (**Fig. S1, Table S1**). Next, we aimed to translate our in vitro observations to live cell imaging, and as such, we fused CLUSTER_277_ to a HaloTag bearing a nuclear localization sequence (NLS) for transfection in HEK293T cells, with HTP–SNAP–NLS (ref^17^) serving as a control (**Fig. 2F**). Indeed, for both cases we observed very good signal overlap from the nuclei, quantified by co-staining (**Fig. S2A**), and SDS-PAGE revealed spliced fragments only in the CLUSTER-expressing cells (**Fig. S2B**). Similar behaviour was observed when we expressed CLUSTER_277_–HTP in the cytosol (**Fig. 2F**).

**Figure 2.**
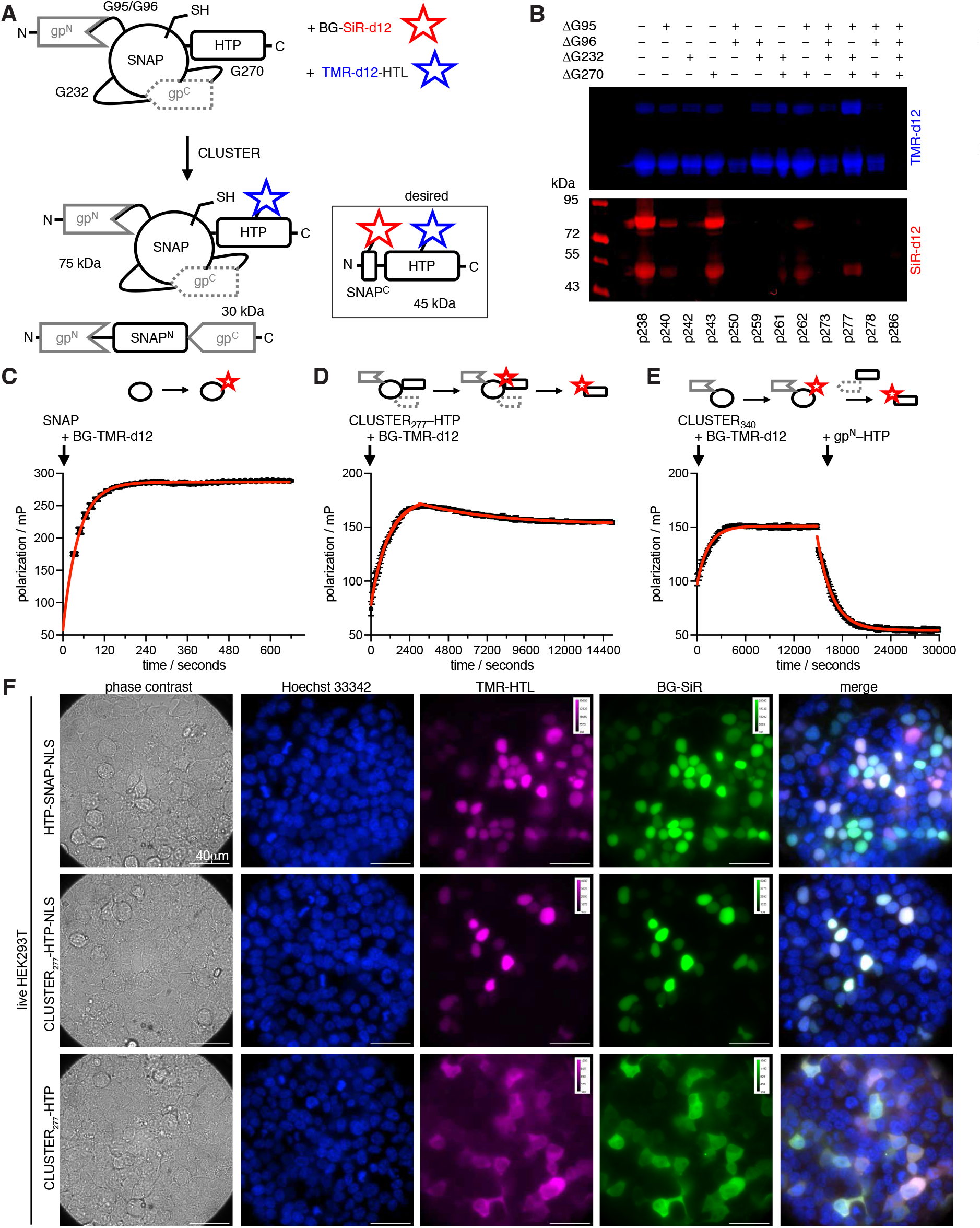
**A)** CLUSTER is equipped with a *C*-terminal HaloTag for orthogonal control, as well as glycine linkers at domain intersections (G95, G96, G232, G270) for subsequent mutational screening. Upon labelling with respective fluorophores, the construct is spliced, releasing the dual-stained SNAP^C^-HTP fragment. **B)** SDS-PAGE gel under a fluorescence imager after CLUSTER-HTP-expression and labelling in *E. coli*. CLUSTER_277_ shows clean labelling of only spliced product. **C-E)** Fluorescence polarization assay to mechanistically determine protein labelling and dynamics. **F)** Live-cell confocal microscopy of HEK293T cells transiently transfected with HTP–SNAP–NLS, CLUSTER_277_–HTP–NLS, and CLUSTER_277_–HTP. Staining was performed overnight using Hoechst 33342, BG-SiR-d12, and TMR-d12-HTL.

To further investigate CLUSTER dynamics, as well as to provide evidence for the anticipated mechanism, the *trans*-splicing variant, CLUSTER_340_, was subjected to a series of computational calculations. The sequential reaction of CLUSTER should consist of 1) labelling reaction with the fluorophore, 2) “unfolding” (or, more general, *destabilization*) of the construct, specifically reducing the rigidity of the SNAP domains to enable association of the intein domains, and 3) splicing, which is conducted via the assembled intein fragments. While both the first and last step of this reaction sequence could be directly measured by time-dependent fluorescence polarization as described above, the second step – the intrinsic destabilization of the construct upon covalent reaction with the fluorophore – remains more speculative than evidence-based. Countering this, the post-labelling (and pre-splicing) stability of CLUSTER_340_ was investigated using molecular dynamics (MD) simulations. For this, the protein structure was predicted using Boltz2^18^ (root mean squared deviation (RMSD) vs. native SNAP-tag, PDB-6Y8P = 0.54 Å, ref^19^), and the ligand moiety of BG-TMR was covalently docked to the active cysteine residue using GNINA 1.3^20^ (**Fig. 3A**). All MD simulations were performed using GROMACS 2025.4 and its included subprograms.^21–31^ After relaxation and equilibration, the unlabelled CLUSTER_340_ protein and the labelled CLUSTER_340_:TMR complex were simulated for 1000 ns in isolation, where the trajectories were produced in triplicates. Alignment of the unwrapped trajectories and analysis of the RMSD of all C-α atoms throughout the trajectories reveals a significantly higher overall perturbation introduced upon covalent binding of TMR (**Fig. 3B**). To further elucidate the spatial destabilization of the protein upon binding, the root mean squared flexibility (RMSF) of the same target atoms was calculated across all recorded MD trajectories, and the difference between the unlabelled and labelled construct was compared (**Fig. 3C**). For a more graphic illustration, the calculated average RMSF values were mapped onto the initially predicted protein structure of CLUSTER_340_ (**Fig. 3D**). Apparently, throughout all recorded trajectories, the unlabelled protein consistently exhibits a higher degree of stability and internal retention of the tertiary structure. Comparison of the RMSD traces reveals a stronger deviation from the equilibrated structure for the labelled construct, which could potentially be associated with stronger drive for spatial rearrangement to properly accommodate the covalently attached ligand. Inspection of the RMSF traces also attributes the arrangements primarily to the residues of the SNAP domains and not the circularly inserted *C*-terminal intein, which further demonstrates the geometric perturbation introduced by the ligand. The insignificant difference in the gp41-1^C^ region could be interpreted as an indicator for the retention of intein function and the potential for post-labelling splicing upon association with the respective *N*-terminal fragment. This association is facilitated and enabled by the increased flexibility of the otherwise rigid SNAP domains after labelling, so that premature splicing is potentially disfavored relative to the labelled construct.

**Figure 3.**
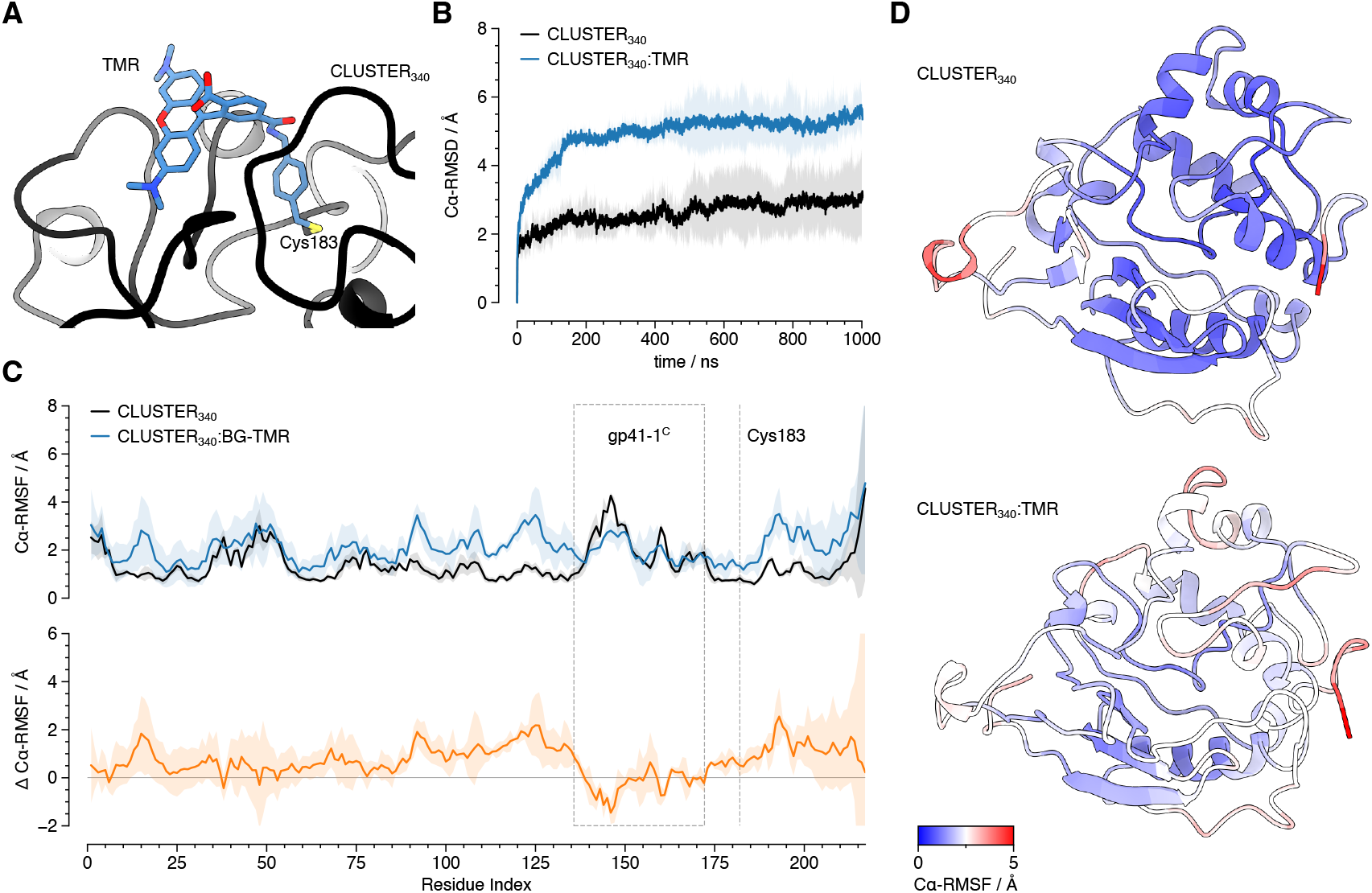
**A)** BG-TMR was covalently docked to the computationally predicted structure of CLUSTER_340_ for subsequent molecular dynamics simulation. **B)** C-α RMSD traces over molecular dynamics simulation trajectories of CLUSTER_340_ and CLUSTER_340_:TMR (opaque: average of triplicate runs, light: absolute spread of measured values across all trajectories). **C)** Top: RMSF of CLUSTER_340_ and CLUSTER_340_:TMR across triplicate molecular dynamics simulation trajectories (opaque: average of triplicate runs, light: absolute spread of measured values across all trajectories). Bottom: Difference of trajectory RMSF between CLUSTER_340_ and CLUSTER_340_:TMR. **D)** Predicted structure of CLUSTER_340_ with color-mapped average RMSF values from **C**.

## DISCUSSION

In summary, this engineered CLUSTER cassette adds a small, fluorescent label at virtually any position (i.e. *N*/*C-*terminally or in between domains) with its strictly stoichiometric reaction. Moreover, the exogenous fluorophore application allows for exceptional temporal control. After splicing, the size (∼5.1 kDa + cargo) of CLUSTER is substantially smaller than the HaloTag Protein (33 kDa), SNAP-tag (20 kDa), BromoCatch (12.8 kDa)^32^ and FPs (27 kDa).^33^ Still bigger than unnatural amino acids,^34^ CLUSTER however overcomes other challenges in terms of costs, toxicity, and putative off-target incorporation.^35^ In contrast, other methods (e.g. Lap-tag/LplA^W37V^ ref.^36^ or TC-tag/ReAsH^37^) offer restricted cargos, or do not covalently react with a dye (e.g. FAST^38^, miniVIPER^39^). While the initial construct made use of a *pseudo-cis* splicing mechanism, leveraging the intramolecular association of originally *trans*-splicing intein domains, a conceptually different approach was demonstrated in CLUSTER_340_. Being originally planned as a diagnostic tool to study the feasibility and discern the contributing reaction rates to the apparent kinetics, CLUSTER_340_ enables true *trans*-splicing of the labelled *C*-terminal SNAP domain onto a POI. By effectively replacing the genetically fused *N*-terminal intein domain with the labelled residue, this strategy mitigates structural perturbations of the POI arising from additional protein bulk. Together, we envision CLUSTER_340_ as a potent labelling agent for biorthogonal and minimally perturbative labelling of accessible protein targets, e.g., cell surface receptors like GLP1R. CLUSTER_277_ demonstrated to function in challenging biological environments like the cytoplasm or inside the mammalian nucleus being fused either *N*- or *C*-terminally. Generally, we envision the utility of CLUSTER to extend to challenging targets and localized spaces, enabling direct observations of small and disordered POIs like tau, synuclein, and GLP-1. The to date limitations remain 1) its comparably slow reaction kinetics (*k*_SNAP_ ∼ 10^4^ M^-1^ s^-1^; *k*_HTP_ ∼ 10^7^ M^-1^ s^-1^) and the 2) not full splicing reaction in mammalian systems. Similar to SNAP- and HaloTags,^40–43^ further protein engineering is needed to address these drawbacks, however, our system demonstrates a plug-and-play approach for proteins by merging a self-labelling tag with intein splicing. We anticipate CLUSTER to further demonstrate superior performance in currently impractical experimental setups, and as such, we envision wide adoption of this method across scales in biomedical and life science applications. In addition, we further anticipate that the reduced steric footprint of CLUSTER, combined with the use of bright synthetic fluorophores, will enable advanced fluorescence-based imaging applications. In particular, these properties may facilitate quantitative single-molecule approaches such as fluorescence correlation spectroscopy (FCS) and fluorescence lifetime imaging microscopy (FLIM), where probe size, concentrations and photophysical stability are critical determinants of performance.^44^ Moreover, the small tag size and compatibility with high-performance dyes are expected to support super-resolution microscopy, potentially improving localization precision and reducing linkage error compared to conventional protein-based labels.^45^

## SUMMARY

In summary, we have designed and cloned a protein chimera consisting of a SNAP-tag in which an *N*-intein was added to its *N*-terminus, while its parent *C*-intein was circularly inserted upstream of the reactive cysteine residue for covalent labelling with BG substrates. While the first system, CLUSTER_238_, was labelled in mammalian cells *N*-terminally on a localized transmembrane protein and C-terminally on a disordered protein (Tau), we optimized the linkers in an *E. coli* screen to yield a variant (CLUSTER_277_) that only showed signals stemming from SNAP-tag substrates in its spliced form. We lastly show the mechanism of reaction by means of fluorescence polarization measurements, including a split version (CLUSTER_340_), for which we lastly performed MD simulations that indicate destabilization after covalent reaction. We anticipate updated CLUSTERs in the future for applications in which small tags are in high demand.

## MATERIALS and METHODS

can be found in the SUPPORTING INFORMATION

## ACKNOWLEDGEMENTS

This project has received funding from the European Union’s Horizon Europe Framework Program (deuterON, grant agreement no. 101042046 to JB). We thank Ramona Birke for technical assistance. Funded by Boehringer Ingelheim Foundation (Exploration Grant to JB). We thank Ines Kretzschmar (Leibniz-FMP) for support on SPPS.

## COMPETING INTERESTS

JB receives licensing revenue from Celtarys Research for provision of chemical probes. MDC and FVB have filed a patent application for the CLUSTER technology. The remaining authors declare no competing interests.

