## Supplemental Information for "A bio-orthogonal and covalent 5 kDa small protein tag"

[1] Leibniz-Forschungsinstitut für Molekulare Pharmakologie, 13125 Berlin, Germany.
[2] Berlin Institute of Medical Systems Biology (BIMSB), Max Delbrück Center for Molecular Medicine, 10115 Berlin, Germany.

[#] Equal contribution
[$] Co-leads
* Correspondence should be addressed to:


### General

Labelling reagents are described in ref[1].

### In Vitro Experiments

#### Cloning and Protein Expression

The initial sequence of CLUSTER containing the circularly permutated SNAP between positions 132/133 to include a split intein $Int^C$ (gp-41-$1^C$) upstream of the reactive Cys145 and a split intein $Int^N$ (gp-41-$1^N$), that resides on the N-terminus of SNAP, was synthesized and purchased from Twist Bioscience HQ (CA, USA). The sequence was inserted into a pET51b(+) vector, replacing SNAP in the original SNAP–Halo–10xHis construct. For all subsequent mutations, respective primer pairs were designed using the NEBaseChanger tool (NewEnglandBiolabs, USA), and site-directed mutagenesis was performed using the NEB Phusion® High-Fidelity DNA Polymerase (NewEnglandBiolabs, USA). Upon successful PCR, the plasmid was transformed into $CaCl_2$ competent *E. coli* cells from the DH5-alpha strain.

After growing a respective liquid bacterial culture for 16 h at 37 °C in LB medium containing 100 µM ampicillin, the medium was removed and plasmid isolation was performed using the NEB Monarch® Plasmid Miniprep Kit (NewEnglandBiolabs, USA). The mutation was confirmed using the Sanger sequencing service offered by LGC Genomics GmbH (Berlin, Germany), whereas the T7 promoter and T7 terminator were used as sequencing primers.

Concluding the bacterial pipeline, the validated plasmid was transformed into $CaCl_2$ competent *E. coli* cells from the BL21(DE3) strain. After growth in LB medium containing 100 µM ampicillin at 37 °C until an $OD_{600}$ of 0.6 was reached, the culture was supplied with 1 µM of IPTG and the temperature was lowered to 18 °C for protein expression overnight. Respective additives, such as BG-SiR-d12 and TMR-d12-HTL, were added directly alongside IPTG.

#### Protein Purification

After protein expression, the bacterial cell pellet was resuspended in 1 mL PBS. While cooling the suspension on ice, the bacterial culture was lysed using pulse sonication for 1 min. The bacterial debris was removed by centrifugation for 5 min at 13,000 rpm and the supernatant, containing soluble components, was subject to further analyses. Purification of proteins containing a 10xHis-tag was performed using the Thermo Scientific HisPur™ Ni-NTA resin. The protein purity was monitored using the protein A280 program of the Thermo Scientific NanoDrop® One Spectrophotometer. After purification, the isolated proteins were stored at 4 °C for further analysis.

#### SDS-PAGE and Fluorescence Gel Imaging

A 1.5 mL microtube was charged with 50 µL of purified protein sample or unpurified bacterial culture (resuspended in PBS), 40 µL of 4x Laemmli Sample Buffer (non-reducing) and 5 µL of dithiothreitol reducing agent. The mixture was incubated at 95 °C for 10 min. Subsequently, 5 µL of the sample mixture was loaded onto a 12 % SDS-PAGE gel, which was prepared using the Roth Rotiphorese® Gel 30 (37.5:1). For qualitative analysis of the bands, the NEB Colour prestained Protein Standard 10-250 kDa was used. After Electrophoresis, the FastGene® Qstain solution was used to visualise protein bands. Additionally, fluorescence images of the fluorescent probes linked to individual proteins were acquired using the FujiFilm FLA-5000 imaging system.

#### Fluorescence Polarization Endpoints

A well of a Corning® 384-well microplate was charged with a 50 µL sample containing 100 µM of the purified protein and 20 µM of fluorescent dye solution in activity buffer (50 mM HEPES, 50 mM NaCl, 1 mM DTT, 1 ng/µL BSA, pH 7.3). Upon addition of the fluorophore, the mixture was incubated for 2 h at ambient conditions. Then, fluorescence polarisation of the sample was measured against respective blank and reference aliquots using a Tecan Spark® microplate reader. All measurements were performed as triplicates.

##### Fluorescence Polarization Kinetics

The purified protein, resuspended in activity buffer (50 mM HEPES, 50 mM NaCl, 1 mM DTT, 1 ng/µL BSA, pH 7.3), was loaded into a Corning® 384-well microplate well (50 µL). Now, using a Tecan Spark® microplate reader, BG-TMR-d12 in activity buffer was injected using the integrated microinjection pump. The final protein concentration was 200 nM, the final fluorophore concentration was 50 nM. After 1 s of orbital shaking, the fluorescence polarization of the sample was measured using excitation at 530 nm and detecting emission using a LP 590 nm filter (30 reads, 40 µs integration time). All measurements were performed at room temperature and in triplicate.

#### Cell Culture and Microscopy

##### Cloning of CLUSTER Constructs for Live Cell Application

The initial sequence of CLUSTER containing the circularly permutated SNAP between positions 132/133 to include a split intein $Int^C$ (gp-41-$1^C$) upstream of the reactive Cys145 and a split intein $Int^N$ (gp-41-$1^N$), that resides on the N-terminus of SNAP, was synthesized and purchased from Twist Bioscience HQ (CA, USA). This sequence was then inserted to SNAP-TM-Halo and TauSNAP vectors, replacing the original SNAP. The TauSNAP vectors were constructed based on the pEGFP-n1-APP vector backbone (Addgene ID69924), including a CMV promotor for protein expression. All plasmids were cloned via Gibson Assembly using NEBuilder HiFi DNA Assembly Master Mix (NewEnglandBiolabs, USA). Primers were designed with NEBuilder Assembly Tool (NewEnglandBiolabs, USA) and listed in table 1. All PCRs were performed with Q5 High Fidelity DNA Polymerase (NewEnglandBiolabs, USA), 10 mM dNTP-Mix (Thermo Fisher Scientific, USA) and all primers were synthesized by Eurofins Genomics Europe (Germany). For site-directed mutagenesis, mutations were inserted using the Q5 Site-Directed Mutagenesis Kit (NewEnglandBiolabs, USA).

##### Cell Culture

HEK293T cells were grown at 37°C under a humified atmosphere of 5% $CO_2$. HEK293T cells were cultured in DMEM/F-12 (Dulbecco´s modified Eagle Medium F-12, no phenol red) plus 10% fetal bovine serum (FBS), 100 U/ml penicillin and 100 µg/ml streptomycin. Cells were passaged upon reaching 90-100% confluence with 0.05% Trypsin-EDTA (Life Technologies, Germany), and used in experiments. Cells used in experiments were pelleted and resuspended in fresh media lacking Trypsin-EDTA.

For imaging of HEK293T cells, cells were grown in eight-well-ibidi treat chambers (µ-slide, eight-well glass bottom, ibidi GmbH, Germany) and for protein lysates in 24-well plates (Sarstedt AG & Co.KG, Germany). Chambers were seeded with 20.000 cells/well and 24-well plates were seeded with 40.000 cells/well and cultured 24h before transfection. HEK293T cell line was confirmed for the absence of mycoplasma using the Venor GeM OneStep mycoplasma detection kit (Minerva Biolabs, Germany).

#### Transfection and Staining

HEK293T cells were transfected with plasmids encoding the Snap-tagged Tau protein or Cluster-tagged Tau protein by X-tremeGENE HP DNA transfection reagent (Merck KGaA, Germany), following the manufacturer´s directions. The HEK293T cells were grown to 70% confluency in the eight-well ibidi chambers (for imaging) and in 24-well plates (for protein lysates) before transfection. Transfections were performed with 250 ng (µ-slide ibidi) and 500 ng (24-well plate) final concentration of plasmids per 250 µl (µ-slide ibidi) and 500 µl (24-well plate) media. Transfection media was removed from cells 24 h later and fresh media was added. For staining, BG-SiR-d12 or TMR-d12-HTL (1 mM stock concentration, 0.5 µM working concentration) and Hoechst 33342 (2 µM working concentration, Merck KGaA, Germany) were added to cells and incubated at 37 °C under a humified atmosphere of 5% $CO_2$ for 30 minutes. For imaging, cells were then washed with fresh media before imaging.

#### Cell Imaging and Analysis

Living cells were imaged in fluorobrite (Invitrogen) using an epifluorescence Nikon Ti-E microscope, equipped with pE4000 (cool LED), Penta Cube (AHF 66-615), 60× oil NA 1.49 (Apo TIRF Nikon) and imaged on a sCMOS camera (Prime 95B, Photometrics) operated by NIS Elements (Nikon). For excitation the following LED wavelengths were used: Hoechst – 405 nm, $JF_{549}$, TMR-d12 – 550 nm, $JF_{646}$, SiR-d12 or on a Leica Stellaris 8-Falcon-STED confocal microscope (Leica Microsystems). To minimize spectral crosstalk between fluorophores, channels were acquired sequentially, starting from the longest to the shortest excitation wavelength. Images were collected at 12-bit depth, and the pixel size was adjusted according to the zoom factor to achieve an optimized spatial resolution. The system is part of the Systems Biology Imaging platform at Max Delbrück Center (MDC), Berlin.

#### Cell Lysis and SDS-PAGE

Transfected HEK293T cells were lysed 24 h after transfection with ice cold RIPA lysis buffer (150 mM NaCl, 10 mM Tris-HCl, 1 mM EDTA, 1% Triton X-100, 0,1% SDS, 0,1% Sodium deoxycholate, complete Protease Inhibitor Cocktail) chilled on ice for 1 h, centrifuged at 13.000 g for 10 minutes at 4 °C. The supernatant was then transferred to a new 1,5 ml reaction tube (Sarstedt AG & Co.KG, Germany) and the protein concentration was detected via Pierce BCA Protein Assay Kit (Thermo Fisher Scientific, USA). For each sample, a total protein concentration of 10 µg was added to 2x loading buffer (50 mM Tris base, 100 mM DTT, 4% SDS, 20% glycerol, bromphenolblue) and a Color Prestained Protein Standard (NewEnglandBiolabs, MA, USA) were loaded on a Bolt 4-12% Bis-Tris Mini-Protein-Gel (Thermo Fisher Scientific, USA). Proteins were separated using a pre-cast, 4%–12% polyacrylamide gel (Thermo Fisher Scientific, USA), at 180 V for 40 minutes in MES SDS running buffer (Thermo Scientific). After electrophoresis, gel was taken from the cast and imaged using a Vilber Fusion FX Imager (Vilber, France) at 680 nm.

#### Statistical Analysis

Data are expressed as the mean±SD and were examined by a one-way analysis of variance ($n$ = 3). More than 3 experiments were performed, and similar results were obtained. $P < 0.05$ was considered to be significant.

### Protein Sequences

SNAP$^{N}$
SNAP$^{C}$
gp41-1$^{N}$
gp41-1$^{C}$

> CLUSTER$_{238}$
MTRSGYCLDLKTQVQTPQGMKEISNIQVGDLVLSNTGYNEVLNVFPKSKKKSYKITLEDGKEIICTEE
HLFPTQTGEMNISGGLKEGMCLYVKE**GG**MDKDCEMKRTTLDSPLGKLELSGCEQGLHRIIFLGKGTSA
ADAVEVPAPAAVLGGPEPLMQATAWLNAYFHQPEAIEEFPVPALHHPVFQQESFTRQVLWKLLKVVKF
GEVISYSHLAALAGNPAATAAVKTALS**G**MMLKKILKIEELDERELIDIEVSGNHLFYANDIL**T**HNG**G**N
PVPILIP**C**HRVVQGDLDVGGYEGGLAVKEWLLAHEGHRLGKR

> CLUSTER$_{dm}$
MTRSGYCLDLKTQVQTPQGMKEISNIQVGDLVLSNTGYNEVLNVFPKSKKKSYKITLEDGKEIICTEE
HLFPTQTGEMNISGGLKEGMCLYVKE**GG**MDKDCEMKRTTLDSPLGKLELSGCEQGLHRIIFLGKGTSA
ADAVEVPAPAAVLGGPEPLMQATAWLNAYFHQPEAIEEFPVPALHHPVFQQESFTRQVLWKLLKVVKF
GEVISYSHLAALAGNPAATAAVKTALS**G**MMLKKILKIEELDERELIDIEVSGNHLFYANDIL**A**HNG**G**N
PVPILIP**C**HRVVQGDLDVGGYEGGLAVKEWLLAHEGHRLGKR

> CLUSTER$_{277}$
MTRSGYCLDLKTQVQTPQGMKEISNIQVGDLVLSNTGYNEVLNVFPKSKKKSYKITLEDGKEIICTEE
HLFPTQTGEMNISGGLKEGMCLYVKE**G**MDKDCEMKRTTLDSPLGKLELSGCEQGLHRIIFLGKGTSAA
DAVEVPAPAAVLGGPEPLMQATAWLNAYFHQPEAIEEFPVPALHHPVFQQESFTRQVLWKLLKVVKFG
EVISYSHLAALAGNPAATAAVKTALSMMLKKILKIEELDERELIDIEVSGNHLFYANDIL**T**HNGNPVP
ILIP**C**HRVVQGDLDVGGYEGGLAVKEWLLAHEGHRLGKR

> CLUSTER$_{340}$
MDKDCEMKRTTLDSPLGKLELSGCEQGLHRIIFLGKGTSAADAVEVPAPAAVLGGPEPLMQATAWLNA
YFHQPEAIEEFPVPALHHPVFQQESFTRQVLWKLLKVVKFGEVISYSHLAALAGNPAATAAVKTALS**G**
MMLKKILKIEELDERELIDIEVSGNHLFYANDIL**T**HNGNPVPILIP**C**HRVVQGDLDVGGYEGGLAVKE
WLLAHEGHRLGKR

> CLUSTER$_{277}$-HTP-10xHis
MASWSHPQFEKGADDDDKVPHMTRSGYCLDLKTQVQTPQGMKEISNIQVGDLVLSNTGYNEVLNVFPK
SKKKSYKITLEDGKEIICTEEHLFPTQTGEMNISGGLKEGMCLYVKE**G**MDKDCEMKRTTLDSPLGKLE
LSGCEQGLHRIIFLGKGTSAADAVEVPAPAAVLGGPEPLMQATAWLNAYFHQPEAIEEFPVPALHHPV
FQQESFTRQVLWKLLKVVKFGEVISYSHLAALAGNPAATAAVKTALSMMLKKILKIEELDERELIDIE
VSGNHLFYANDIL**T**HNGNPVPILIP**C**HRVVQGDLDVGGYEGGLAVKEWLLAHEGHRLGKRGRLEVLFQ
GPKAFLEGSEIGTGFPFDPHYVEVLGERMHYVDVGPRDGTPVLFLHGNPTSSYVWRNIIPHVAPTHRC
IAPDLIGMGKSDKPDLGYFFDDHVRFMDAFIEALGLEEVVLVIHDWGSALGFHWAKRNPERVKGIAFM
EFIRPIPTWDEWPEFARETFQAFRTTDVGRKLIIDQNVFIEGTLPMGVVRPLTEVEMDHYREPFLNPV
DREPLWRFPNELPIAGEPANIVALVEEYMDWLHQSPVPKLLFWGTPGVLIPPAEAARLAKSLPNCKAV
DIGPGLNLLQEDNPDLIGSEIARWLSTLEISGAPGFSSISAHHHHHHHHHH

> SNAP$^{C}$ (from SPPS)
**T**HNGGNPVPILIP**C**HRVVQGDLDVGGYEGGLAVKEWLLAHEGHRLGKR

### Molecular Modelling

#### Structure Preparation and Covalent Docking

The initial receptor structure was predicted from the amino acid sequence using Boltz2. Structure prediction was performed with multiple sequence alignment support and potential-based refinement enabled. The resulting structural models were protonated for physiological conditions using OpenBabel 3.1 and used directly for subsequent docking.

The ligand structure was generated from SMILES representation using OpenBabel 3.1 and converted to three-dimensional geometry with standard protonation at physiological conditions and using the conformer generation setting “Best”. The structure was saved in SDF format and used without further modification.

Covalent docking was performed using GNINA 1.3. The predicted protein structure was used as the receptor, and the docking search space was defined automatically using the full receptor structure with additional padding of 5.0 Å and automatic box extension enabled. Covalent attachment was specified between the sulfur atom of residue Cys183 and the respective benzylic carbon atom as a single bond. Flexibility of the receptor was introduced locally by allowing side-chain flexibility for the covalent cysteine.

Docking was performed using the Vinardo scoring function, followed by convolutional neural network (CNN)-based rescoring with the general_default2018 parametrization. The search exhaustiveness was set to 128, and the random seed was set to 100. The docked poses were ranked by the CNN confidence score, and the highest scoring complex was used as starting point for subsequent molecular dynamics simulations.

#### Molecular Dynamics Simulation

All molecular dynamics simulations were performed using GROMACS 2025.4. First, the protein structure was protonated for physiological conditions using PDBFixer, a tool supplied within the OpenMM 8.2 suite. Subsequently, the protein was parametrized using the AMBER14SB protein force field including improper dihedrals, as supplied in the OpenFF toolbox. All non-standard bonds and atoms were parametrized using the OpenFF Sage 2.3 force field. The system was solvated in a dodecahedral box of TIP3P water with a minimum solute-box distance of 10 Å and neutralized with NaCl to a final concentration of 150 mM.

Periodic boundary conditions were applied in all dimensions. Short-range electrostatic and van-der-Waals interactions were truncated at 1.0 nm, and long-range electrostatics were treated using the particle mesh Ewald (PME) method. Neighbour searching was performed using the Verlet cutoff scheme with an update frequency of 20 steps. Analytical dispersion corrections to energy and pressure were applied.

Energy minimization was carried out using the steepest descent algorithm for up to 50,000 steps with an initial step size of 0.01 nm and a convergence cutoff of 500.0 kJ $mol^{-1}$ $nm^{-1}$. The minimized system was first equilibrated in the NVT ensemble for 5 ns at 300 K using a 1 fs integration time step. Temperature coupling was achieved using the velocity-rescaling thermostat (V-rescale) with a coupling constant of 0.1 ps. Subsequently, the systems were equilibrated in the NPT ensemble for 5 ns at 300 K and 1 bar using the Parinello-Rahman barostat (coupling constant of 2.0 ps, compressibility 4.5e-5 $bar^{-1}$).

All bonds involving hydrogen atoms were constrained using the LINCS algorithm (order 4, one iteration). Production simulations were then performed in the NPT ensemble for 1000 ns using a 2 fs integration time step under identical thermostat and barostat settings. All simulations were run as triplicates. Coordinates were written every 100 ps during production, unwrapping and alignment of the trajectories was performed using GROMACS.

### Supplementary Figures

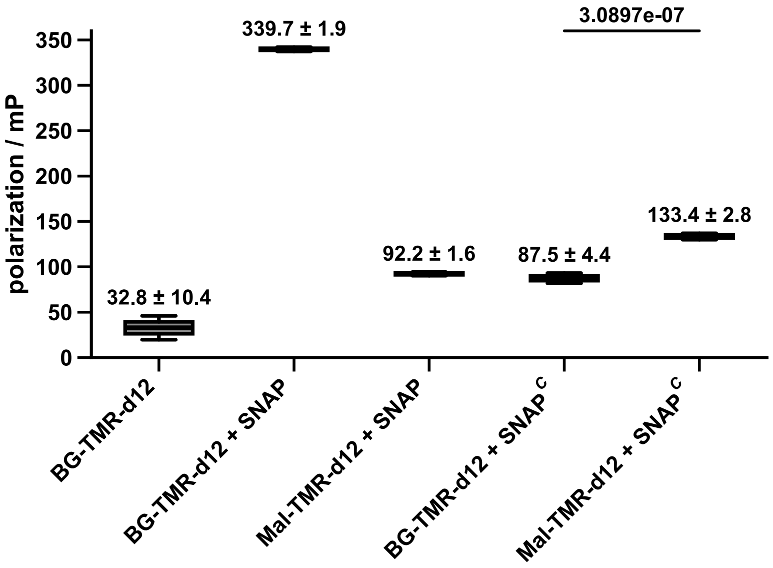

**Figure S1. Fluorescence polarization measurements at reaction endpoints.** The purified protein/peptide was incubated with the respective substrate for 2 h in activity buffer before fluorescence polarization was measured. All measurements were performed at room temperature, *n* = 5, reported values are mean ± SD**.** Unpaired t-test, *p* = 3.0897e-7.

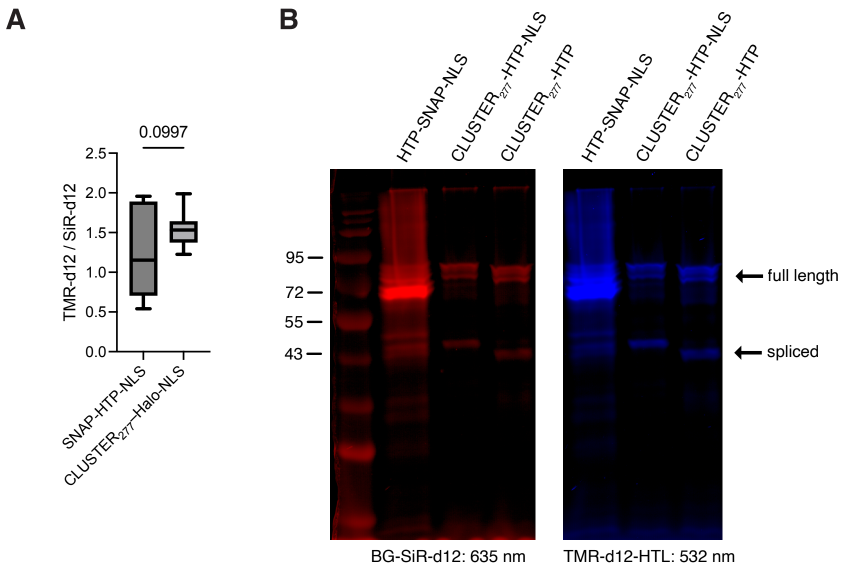

**Figure S2. Additional data from Figure 2. A)** Integrated intensity of TMR-d12 divided by SiR-d12 channel for HTP-SNAP-NLS and CLUSTER$_{277}$-HTP-NLS transfected cells. n = 10 ROIs, Unpaired t-test, p=0.0997. **B)** Fluorescent image of the SDS-PAGE of cell lysates from HTP-SNAP-NLS, CLUSTER$_{277}$-HTP-NLS and CLUSTER$_{277}$-HTP transfected HEK293T cells.

### Supplementary Tables

**Table S1. Mass spectrometry of SNAP$^C$ ± ligands.**

| | expected (Da) | observed (Da) |
|---|---|---|
| SNAP$^C$ | 5166 | 5165 |
| SNAP$^C$ + Mal-TMR-d12 | 5731 | 5731 |
| SNAP$^C$ + BG-TMR-d12 | 5710 | 5165 |
